# Evolutionary history of MEK1 illuminates the nature of cancer and RASopathy mutations

**DOI:** 10.1101/2023.03.09.531944

**Authors:** Ekaterina P. Andrianova, Robert A. Marmion, Stanislav Y. Shvartsman, Igor B. Zhulin

## Abstract

Mutations in signal transduction pathways lead to various diseases including cancers. MEK1 kinase, encoded by the human *MAP2K1* gene, is one of the central components of the MAPK pathway and more than a hundred somatic mutations in *MAP2K1* gene were identified in various tumors. Germline mutations deregulating MEK1 also lead to congenital abnormalities, such as the Cardiofaciocutaneous Syndrome and Arteriovenous Malformation. Evaluating variants associated with a disease is a challenge and computational genomic approaches aid in this process. Establishing evolutionary history of a gene improves computational prediction of disease-causing mutations; however, the evolutionary history of MEK1 is not well understood. Here, by revealing a precise evolutionary history of MEK1 we construct a well-defined dataset of MEK1 metazoan orthologs, which provides sufficient depth to distinguish between conserved and variable amino acid positions. We used this dataset to match known and predicted disease-causing and benign mutations to evolutionary changes observed in corresponding amino acid positions. We found that all known and the vast majority of suspected disease-causing mutations are evolutionarily intolerable. We selected several MEK1 mutations that cannot be unambiguously assessed by automated variant prediction tools, but that are confidently identified as evolutionary intolerant and thus “damaging” by our approach, for experimental validation in *Drosophila*. In all cases, evolutionary intolerant variants caused increased mortality and severe defects in fruit fly embryos confirming their damaging nature predicted by out computational strategy. We anticipate that our analysis will serve as a blueprint to help evaluate known and novel missense variants in MEK1 and that our approach will contribute to improving automated tools for disease-associated variant interpretation.

**Significance Statement:** High-throughput genome sequencing has significantly improved diagnosis, management, and treatment of genetic diseases and cancers. However, in addition to its indisputable utility, genome sequencing produces many variants that cannot be easily interpreted – so called variants of uncertain significance (VUS). Various automated bioinformatics tools can help predicting functional consequences of VUS, but their accuracy is relatively low. Here, by tracing precise evolutionary history of each amino acid position in MEK1 kinase, mutations in which cause neurodegenerative diseases and cancer in humans, we can establish whether VUS seen in humans are evolutionarily tolerant. Using published data and newly performed experiments in an animal model, we show that evolutionarily tolerable variants in MEK1 are benign, whereas intolerable substitutions are damaging. Our approach will help in diagnostics of MEK1-associated diseases, it is generalizable to many other disease-associated genes, and it can help improving automated predictors.

## Introduction

In the last few decades gene sequencing has demonstrated its great potential for diagnostics of inherited disorders and common pathologies in clinical practices. Nowadays, genetic testing is broadly and routinely utilized by researchers and clinicians to identify genetic variations related to the disease, and in many cases helps choosing the therapeutic strategy. However, only a small fraction of reported variants is tested experimentally to establish causality and most newly discovered coding variations are neither described in other individuals nor studied in cellular or animal models. The growing sequencing data has led to massive accumulation of variants of uncertain significance (VUS): currently over 500 000 of reported variants are classified as a VUS in ClinVar, the largest public archive on human genetic variation and phenotypes (1). The lack of experimental data on VUS forces clinicians to rely on predictions using computational tools, such as PolyPhen-2 (2) and SIFT (3); however, the accuracy of these tools is far from perfect. For example, a study that calculated prediction accuracy by comparing predictions for mutation outcome made by SIFT and PolyPhen-2 with the existing experimental data, found that false negative predictions comprised 43% and 33%, respectively (4). Similarly, a large comparative study assessed the performance of ten prediction tools and found that specificities of variant interpretation were 63-67% for SIFT and 73-75% for PolyPhen-2 (5). In some cases, well studied pathogenic mutations were falsely declared to be benign by both tools (6). One of the possible reasons for the erroneous predictions is the inability of the automated algorithms to exclude functionally unrelated homologs from the multiple sequence alignment (MSA) used for variant interpretation. The majority of disease genes have been duplicated in their evolutionary history and usually only one copy of the gene is associated with a disease (7). Therefore, understanding gene’s history and selecting only well-defined orthologs (by excluding paralogs and undefined homologs) for variant interpretation is a critical step in improving evolutionary-based prediction methods.

The mitogen-activated protein kinase kinase 1 (MKK1, MEK1 or MAP2K1) is a member of the vast kinase family and the centerpiece of Ras/MAPK pathway, which plays a vital role in the regulation of cell growth and proliferation. Dysregulation of the Ras/MAPK cascade through activating mutations results in the abnormal signaling, which consequently leads to tumorigenesis and developmental disorders (8-10). Early approaches to simultaneous sequencing of multiple genes of Ras/MAPK signaling pathways have yielded hundreds of mutations in MEK1 possibly associated with RASopathies and cancer. For example, germline MEK1 mutations have been reported in patients with cardiofaciocutaneous (CFC) syndrome (11-13), and somatic mutations have been identified in melanoma (14-16), lung cancer (17, 18), gastric cancer (19), colon carcinoma (20), ovarian cancer (21), hairy cell leukemia (22), Langerhans Cell Histiocytosis (LCH) and the non-LCH (23-25). Due to the central role of MEK1 in the Ras/MAPK pathway signaling, it is a common target for highly selective inhibitors, such as Cobimetinib and Trametinib, that are used in cancer treatment (26-30). MEK inhibition has shown to be very successful in treating various cancers and is now also gaining recognition for treating RASopathies (31).

Seven MEK paralogs are encoded in the human genome (Fig 1). A previous study identified potential duplication events and relationship between the MEK paralogs (32); however, due to a small number of genomes available at the time, that study provided limited information. Here, we reveal a precise evolutionary history of the MEK family and confidently identify MEK1 orthologs in hundreds of metazoan genomes. We generated their curated multiple sequence alignments suitable for interpretation of genetic variants in humans. This allowed us to demonstrate that all known and the vast majority of suspected disease-causing mutations in MEK1 are evolutionarily intolerable. Finally, we selected several variants that could not be unambiguously interpreted by existing prediction tools for experimental validation and show that evolutionary computationally predicted intolerant variants cause increased lethality and severe defects in fruit fly embryos. Based on this, we argue that our computational evolutionary approach is applicable for interpretation of selected VUS in MEK1 and many other human genes.

**Figure 1.**
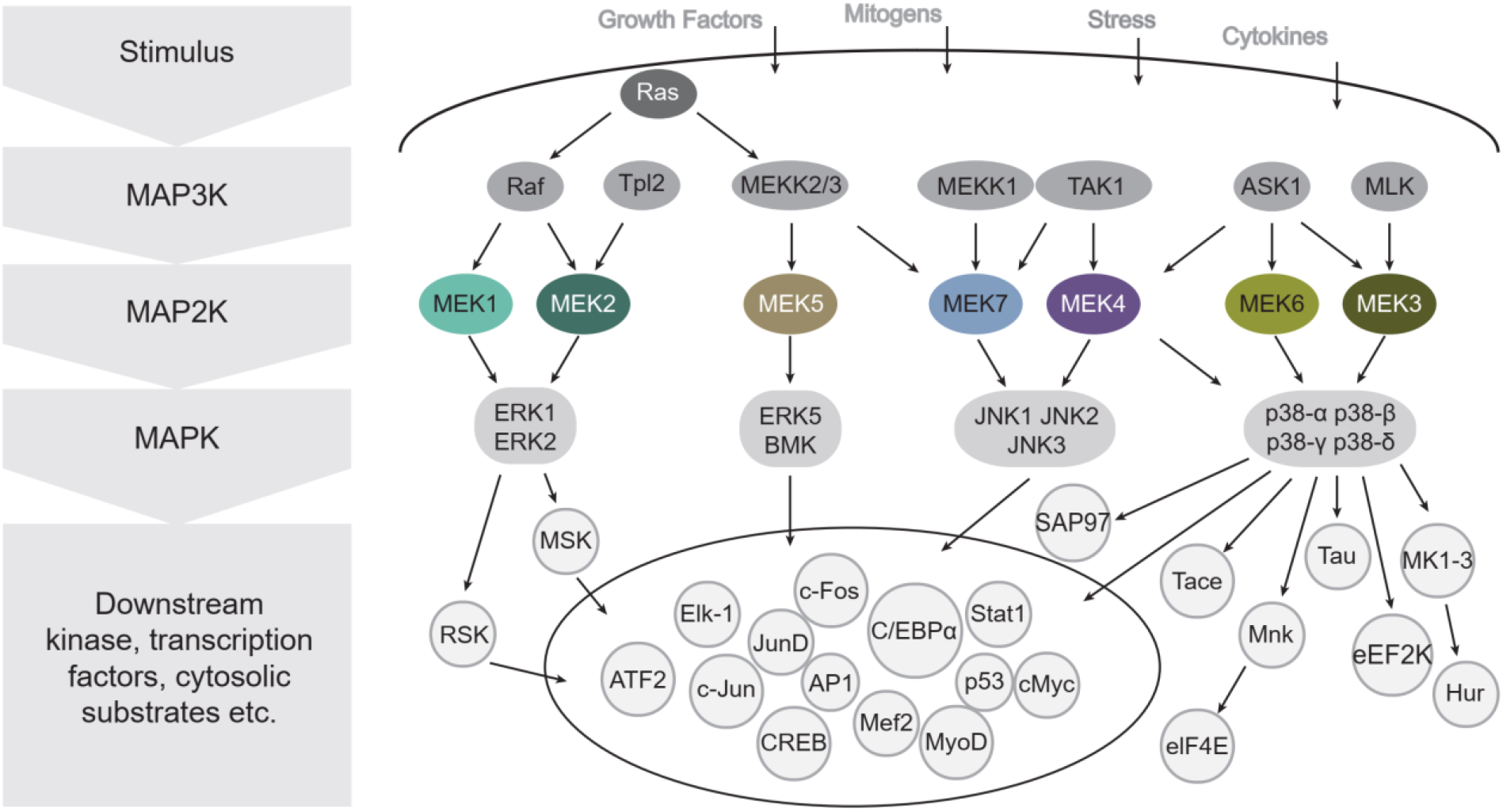
Central role of MAP2K kinases in MAPK/ERK pathway. Arrows indicate known interactions. Adopted from (76)

## Results

### Origins and evolution of the MEK family

To reveal the precise evolutionary history of MEK, we first identified all MEK homologs and their closest relatives within the previously defined STE group of protein kinases (33). We used BLAST searches initiated with human MEK sequences against the selected genome dataset representing all major Eukarya lineages. All sequences collected in these searches were used to generate a multiple sequence alignment that was edited to correct misaligned regions (Dataset S1). The alignment was then used for phylogenetic reconstruction by maximum likelihood and neighbor-joining methods (Fig. 2, Fig. S1 and Fig. S2). Both approaches yielded trees with similar topology, indicating the reliability of phylogenetic inference.

**Figure 2.**
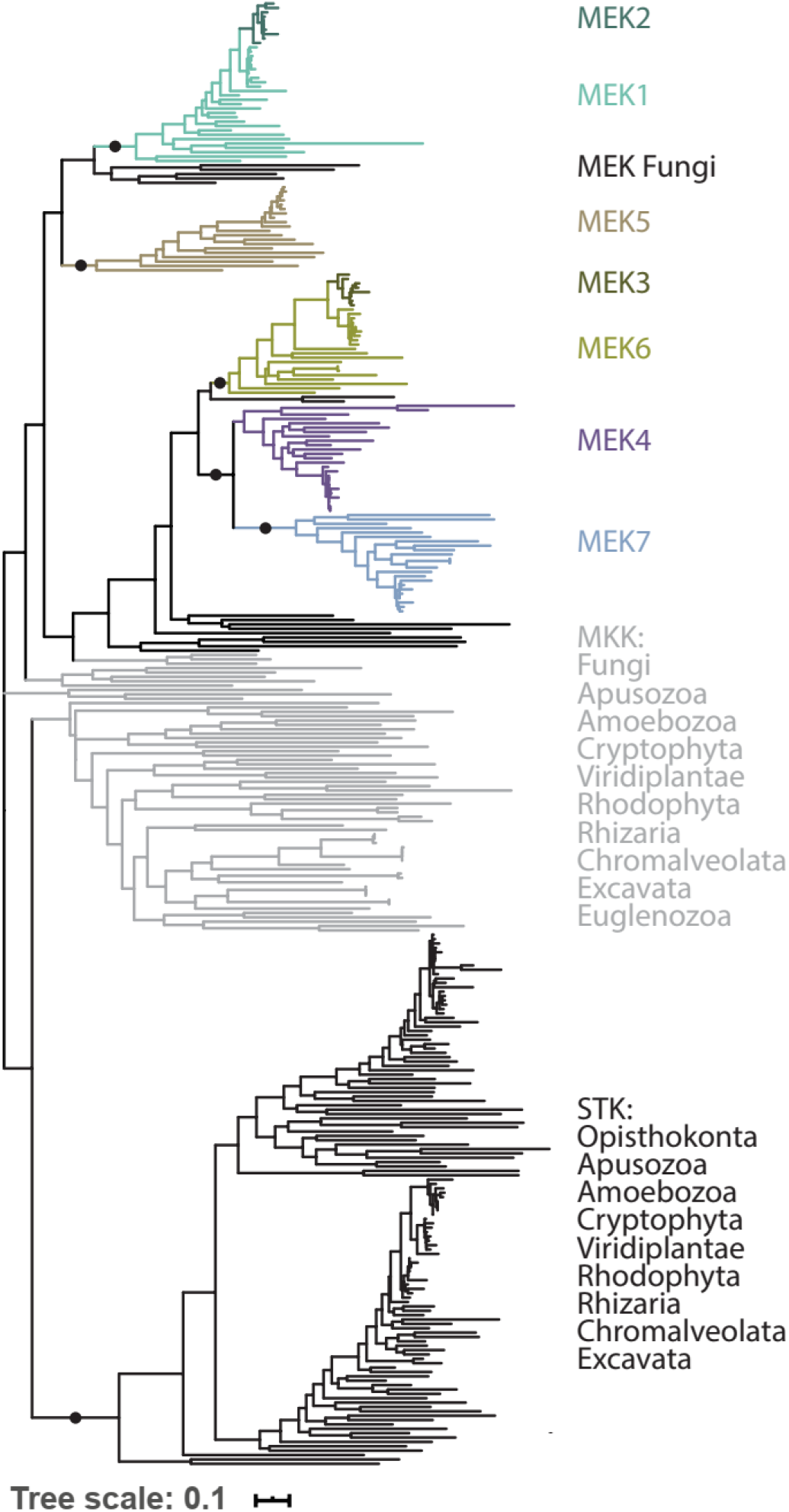
Maximum likelihood phylogenetic tree of the MEK protein family and their closest homologs from the representative set of eukaryotic species. Bootstrap support values >50% are indicated as circles.

The phylogenetic reconstruction revealed three distinct clusters in the representative genome dataset (Fig. 2). The first cluster contained human MST1 (STK3), MST2 (STK4), MST4 (STK26), and YSK1 (STK25) kinases and had at least one sequence from each representative genome from our set. All seven human MEK sequences were found in the second cluster (termed MEK), which contained sequences only from Obazoa. The third cluster (termed MKK) contained sequences from all other major eukaryotic supergroups, including known MKK kinases from *Arabidopsis thaliana*. This cluster was not supported by appreciable bootstrap values; however, the Cluster of Orthologous Groups approach suggested that sequences from MKK cluster are more related to MEK1 than to any other MEK or MST sequences. Within the MEK cluster, sequences were further branching out to form three clades, each containing one or more human MEK proteins, in a good agreement with the early analysis (32). Clade I contained sequences from all Opisthokonta and included human MEK1 and MEK2. Clade II contained sequences from Metazoa, Choanoflagellida and Filasteria and included human MEK5. Larger clade III was almost exclusively composed of sequences from Metazoa and included human MEK4, MEK7, MEK3 and MEK6. The bootstrap values supporting sequences from Choanoflagellida and Filasteria in clade III were weak and, therefore, to validate their orthologous relationships we employed the Cluster of Orthologous Groups approach and assessed sequence and domain synapomorphies. As a result, we assigned each sequence to a respective MEK group and annotated each of the protein sequences from the representative genome set according to the *H. sapiens* MEK nomenclature. Phyletic distribution of MEK homologs (Fig. 3) also suggests that MEK1 is the ancestor of the entire MEK family, which is further supported by the fact that the MEK1 clade within Opisthokonta has the shortest average brunch length from the root (Fig. 2), indicating the least divergence, which is typical of the ancestor. Based on these observations and the presence of MEK homologs in Viridiplantae, Chromalveolata, Excavata, and Rhizaria, we further suggest that MEK1 was present in the last eukaryotic common ancestor and that its separation from MST-type serine/threonine kinases occurred prior to the emergence of major eukaryotic supergroups.

**Figure 3.**
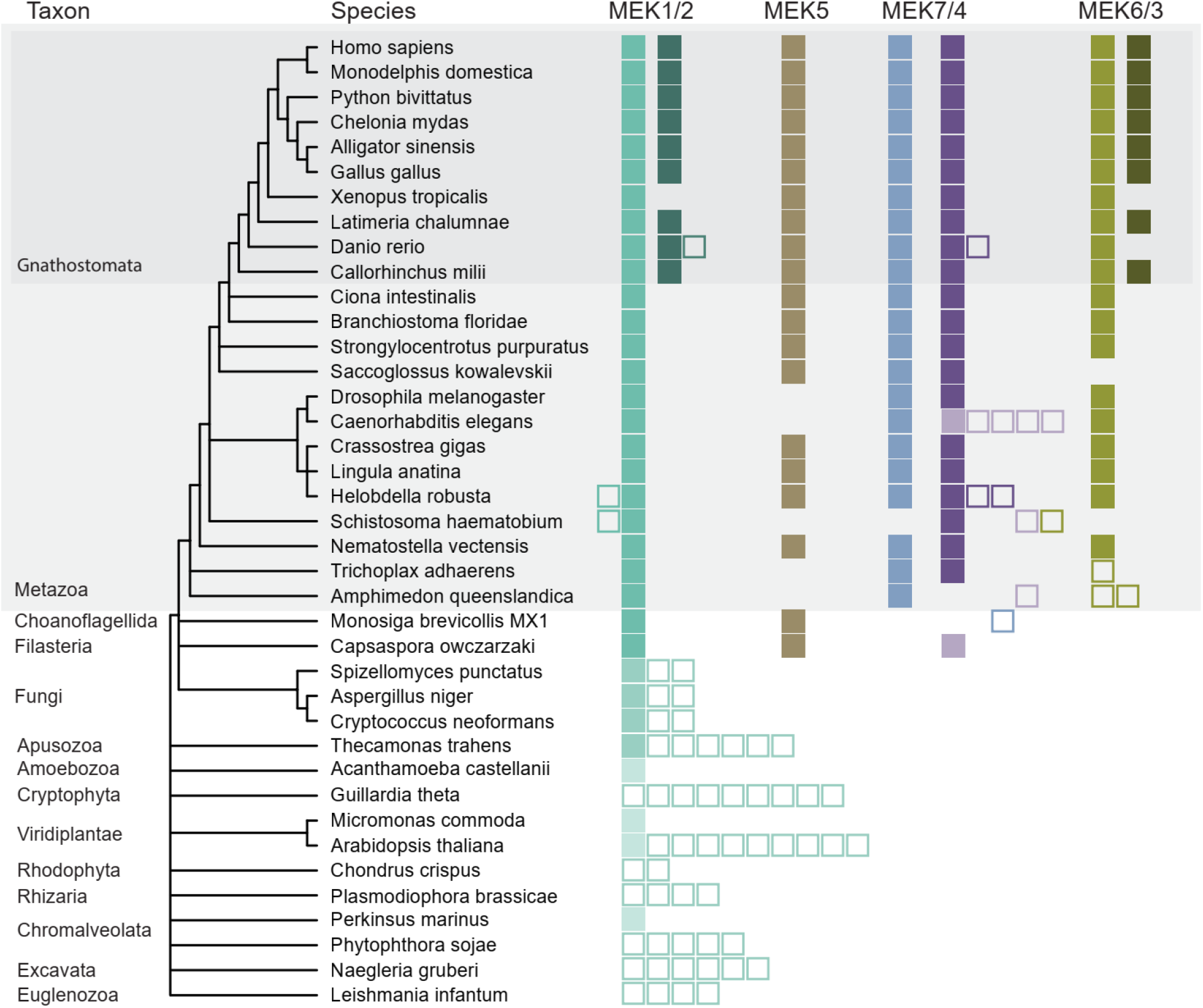
Evolutionary history of MEK family across the major eukaryotic lineages. Schematic eukaryotic tree of life was extracted from NCBI Taxonomy Browser. Each member of MEK family (MEK1 through MEK7) is indicated by a different color. Filled squares represent confidently assigned orthologs. Empty squares represent paralogs (placed next to filled square of corresponding orthologs) or proteins that could not be confidently assigned to a specific MEK class.

Several major duplication events have contributed to diversification of the MEK family in the lineage leading to humans. The first two duplication events resulting in the emergence of MEK5 and the common ancestor of MEK4/7/6/3 most likely have occurred in the last common ancestor of Holozoa, as both *M. brevicollis* and *C. owczarzaki* have homologs that cluster with the corresponding clades. Another major duplication event has occurred at the root of Metazoan lineage, giving rise to MEK6, MEK7 and MEK4. Finally, with the emergence of vertebrates, the MEK family experienced an expansion again, with MEK2 and MEK3 diverging from MEK1 and MEK6, respectively.

### Building a comprehensive set of well-defined MEK1 orthologs

The list of MEK1 orthologs generated from the representative set of Eukarya genomes was used as a starting point and a reference, with the idea of its expansion by “filling the blanks”, *e*.*g*. identifying MEK1 in genomes that were not represented in this dataset. We limited our searches to Metazoan genomes for several reasons. First, our phylogenetic analysis demonstrated that we could confidently distinguish MEK1 orthologs from paralogs in Metazoa. Second, more than two hundred metazoan genomes were available for analysis providing substantial depth. Finally, estimated time since Metazoan emergence, 635 Mya (34) should result in substantial sequence diversity, which is essential for productive sequence analysis and variant interpretation. To generate a comprehensive list of MEK1 orthologs, we searched all metazoan genomes available in RefSeq by BLAST using the human MEK1 sequence as a query. Best BLAST hits from each genome were collected and combined with MEK1 orthologs from the representative genome set generated in the previous step (Metazoa only). MEK2, MEK5, MEK3, MEK6, MEK4 and MEK7 sequences from the same representative set of metazoan genomes were added to this dataset to be used as outgroups in a downstream phylogenetic analysis and all full-length sequences were aligned. Neighbor-joining and maximum likelihood phylogenetic trees were built from the resulting MSA. Based on the phylogenetic inference, outgroups containing MEK2 through MEK7 and several newly identified paralogs were discarded, thus resulting in a master MSA containing more than 300 well-defined MEK1 orthologs from Metazoa (Dataset S2).

### Dataset of confidently assigned MEK1 orthologs provides a necessary and sufficient evolutionary depth

Functionally important positions in proteins are conserved, whereas multiple substitutions observed in a group of orthologs indicate that protein tolerates changes in a given position without losing its function. An informative MSA should contain both conserved and variable positions, indicating that a given set of sequences has a sufficient evolutionary depth. To test our MEK1 ortholog MSA (Dataset S2), we analyzed 323 amino acid positions of the MEK1. We found that 66 amino acid positions (20% of the total analyzed residues) were invariable indicating their critical importance (Dataset S3). Because no changes in these positions were tolerated during more than 600 Mya of MEK1 evolutionary history, any substitution in invariable positions should be considered deleterious. Among invariable positions, 27 were specifically conserved in MEK1 orthologs (variable in other MEK family members): these positions likely define MEK1-specific functions, such as interaction with its cognate partners within the MAPK signaling pathway. We also identified 28 positions, where a single substitution event has occurred (either in a single organism or in a common ancestor of a closely related group). Considering possibilities of a sequence error or a compensatory mutation, confident predictions for such positions should not be made. Finally, the majority of positions (∼70%) were variable (>2 independent substitution events during the evolutionary history). Fig. 4 shows examples of both invariable and highly variable positions in MEK1. For example, valine in position

**Figure 4.**
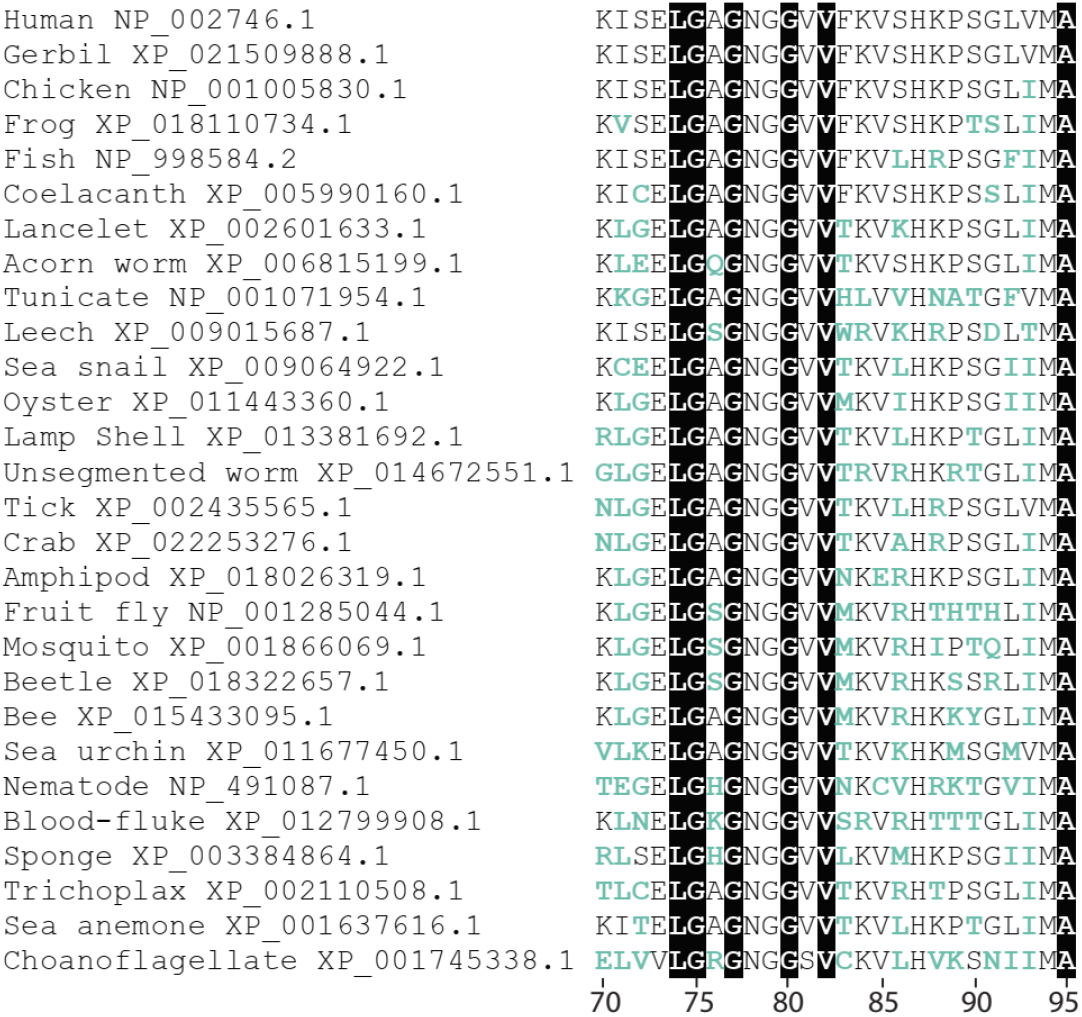
Conserved and variable positions in MEK1 proteins from various species. Small subset of more than 300 MAP2K1 orthologs is shown. Positions highlighted in black are invariable in the entire dataset of MEK1 orthologs. Variable positions are shown in teal.

82 is invariable in the entire dataset of more than 300 sequences, but the neighboring position 83 (phenylalanine in human MEK1) is highly variable: it can be occupied not only by a large hydrophobic residue, such as tryptophan and leucine, but also by small polar residues, such as serine, threonine, and asparagine. Hence, one can conclude that orthologs had enough time to diverge, while preserving key positions, and therefore the generated MEK1 ortholog MSA has necessary and sufficient evolutionary depth to predict consequences of missense mutations in the human MEK1 gene. For example, any change in position V82 in humans (including changes to any similar aliphatic amino acid) would be classified as evolutionarily intolerable and thus damaging. On the other hand, mutation F83S would be classified as tolerable and thus likely benign, even though a large aromatic residue is substituted with a small polar one.

### Functionally important residues are conserved in MEK1 ortholog MSA

Functionally important amino acid residues in proteins, such as catalytic sites, are well conserved. Thus, to validate our MEK1 ortholog MSA, we examined conservation of functionally characterized amino acid residues in MEK1 that are available in the literature (Table S1). As expected, amino acid residues that are required for the kinase catalytic activity or define its key structural properties are conserved not only in the MEK1 ortholog MSA (Dataset S2), but also in the entire MEK homolog MSA (Dataset S1). For example, catalytic core residues K97, D190, and D208 (35), ATP-phosphate binding loop residues G75, G77, and G80 22177953 (35), and catalytic loop residues R189 and K192 (35) are invariable in both datasets. Amino acid residues comprising protein-protein interaction interfaces are usually conserved, but not as strictly as active or catalytic sites. It is worth noting that L314, which participates in MEK1-BRAF and MEK1-KSR1 complex formation (36, 37) is invariable in MEK1 orthologs, thus highlighting its functional importance. Another functionally important residue, E138, forms a salt bridge with BRAF R462 in the heterodimer complex (37). We observe alternations in this position in some MEK1 orthologs (Dataset S2): however, all changes are to polar residues only (aspartate, asparagine, and serine). Thus, interaction between these MEK1 and BRAF residues is preserved: either as a salt bridge or as a weaker hydrogen bond. Taken together, these observations further support the suitability of our MEK1 ortholog MSA for the variant interpretation.

### Known pathogenic mutations are evolutionarily intolerable

We further validated our MEK1 ortholog MSA against available experimental and clinical data. All known disease-causing MEK1 mutations were identified in the literature and are listed in Table 1. Satisfactorily, we found that all experimentally confirmed pathogenic mutations were predicted as such by our approach (Table 1). For example, in position 53 only one type of substitution (F53W, found exclusively in Nematoda) is observed in our MEK1 ortholog MSA (Dataset S2). Therefore, all other substitutions, including well documented F53L and F53S mutations (11, 15) are evolutionarily intolerable. Another aromatic residue, Y130, had changed multiple times during MEK1 evolution, but the type of substitution was always the same - Y130F (Dataset S2). Thus, we can conclude that all other types of substitutions in this position, including a well characterized, disease-causing Y130C mutation (11) are intolerable. Evolutionarily intolerable positions reduce fitness and thus should be defined as “damaging”, “deleterious” and, when the change in protein function leads to a disease, “disease-causing”.

**Table 1.** Experimentally confirmed disease-causing missense mutations in MEK1 are evolutionarily intolerable.

| **Mutation** | **Zygosity** | **Experimental data** | **Association** | **Evolutionary tolerance** |
| --- | --- | --- | --- | --- |
| L42F | heterozygous | phenotypic changes of zebrafish embryos (52) | CFC (13) | No |
| R47Q | homozygous | stimulated ERK phosphorylation, transforming ability (19) | Cancer (19) | No |
| R49L | heterogeneous | stimulated ERK phosphorylation, transforming ability (19) | Cancer (19) | No |
| F53S | heterogeneous | stimulated ERK phosphorylation (11, 85), phenotypic changes of zebrafish embryos (52) | CFC (11) | No |
| F53L | heterozygous | stimulated ERK phosphorylation (15); phenotypic changes of zebrafish embryos (52) | Cancer (15) | No |
| T55P | ND | stimulated ERK phosphorylation (85); phenotypic changes of zebrafish embryos (52) | CFC (47) | No |
| K57N | heterozygous | stimulated ERK phosphorylation (85), transforming ability (59) | Cancer (59) | No |
| D67N | heterozygous | stimulated ERK phosphorylation (21, 85); phenotypic changes of zebrafish embryos (52) | Cancer (21) | No |
| C121S |  | stimulated ERK phosphorylation (85, 86) | Cancer (86) | No |
| P124L | ND | phenotypic changes of zebrafish embryos (52) | CFC (87) | No |
| P124Q | heterozygous | phenotypic changes of zebrafish embryos (52) | Costello syndrome (88) | No |
| P124S | heterozygous | stimulated ERK phosphorylation (15) | Cancer (15) | No |
| G128V | ND | phenotypic changes of zebrafish embryos (52) | CFC (89) | No |
| Y130C | heterozygous | stimulated ERK phosphorylation (11, 85); phenotypic changes of zebrafish embryos (52) | CFC (11) | No |
| Y130H | heterozygous | phenotypic changes of zebrafish embryos (52) | CFC (13) | No |
| Y130N | ND | phenotypic changes of zebrafish embryos (52) | CFC (90) | No |
| E203K | heterozygous/<br>homozygous | stimulated ERK phosphorylation (15, 85); phenotypic changes of zebrafish embryos (52) | Cancer (15) | No |
| E203Q | ND | phenotypic changes of zebrafish embryos (52) | CFC (91) | No |
| I204T | heterozygous | stimulated ERK phosphorylation, transforming ability (19) | Cancer (19) | No |

### Known likely benign mutations are evolutionarily tolerable

MEK1 alleles found in high frequencies in human populations (very recent evolutionary changes) are likely to be benign. Therefore, to use it as a negative control for our interpretation of missense mutations, we compiled a list of MEK1 missense variants with frequency >5 (Table 2) from more than 140,000 human exomes and genomes available in the Genome Aggregation Database (gnomAD, (38)). Remarkably, all these variants were found to be evolutionarily tolerable and therefore predicted as benign. For example, A390T variant is found at high frequency in human cohorts and we identified exactly the same variant in other mammals (*Odobenus rosmarus divergens* and *Sorex araneus*). Furthermore, other substitutions for small amino acids in this position can be seen throughout the evolutionary history of MEK1 (e.g. valine, serine, glycine), reinforcing the notion that the observed A390T variant in humans is benign.

**Table 2.** High frequency MEK1 variants found in human cohorts.

| **Variant** | **gnomAD**<br>**version** | **Allele**<br>**Count** | **Allele**<br>**Frequency** | **Evolutionary changes\*** | **Evolutionary**<br>**tolerance** |
| --- | --- | --- | --- | --- | --- |
| A19G | v3 | 8 | 5.6E-05 | Gly, >2 events | Yes |
| V93I | v2.1.1 | 11 | 3.9E-05 | Ile, > 2 events | Yes |
| A283V** | v3 | 40 | 2,8E-04 | Val, > 2 events | Yes |
| P321S | v2.1.1 | 14 | 4.9E-05 | Ser, 1 event; multiple events of<br>change to other polar aa | Yes |
| N345S | v2.1.1 | 7 | 2.8E-05 | Ser, 1 sequence; multiple events of<br>change to other polar aa | Yes |
| A347T | v2.1.1 | 7 | 2.8E-05 | Thr, >2 events | Yes |
| G354H | v2.1.1 | 11 | 3.9E-05 | His, 1 sequence; multiple events of<br>change to other aa | Yes |
| L375I | v2.1.1 | 11 | 4.4E-05 | Ile, > 2 events | Yes |
| G380S | v2.1.1 | 5 | 2.0E-05 | Ser, 1 event; multiple events of<br>change to other polar aa | Yes |
| A390T | v2.1.1 | 19 | 6.7E-05 | Thr, >2 events | Yes |
| V393I | v2.1.1 | 8 | 2.8E-05 | Ile, > 2 events | Yes |

### MEK1 missense mutations associated with RASopathies and cancer are evolutionarily intolerable

Extensive search through COSMIC and ClinVar databases allowed us to retrieve all missense mutations in MEK1 that were reported to be associated with cancer and/or RASopathy. After excluding repetitive reports, we identified a total of 256 potentially pathogenic variants in the MEK1 protein (Table S2). Among more than a hundred of disease-associated variants, seventeen are found in positions that are invariable (100% identity throughout the MEK1 MSA): any change in any of these positions is evolutionarily intolerable and therefore likely to substantially alter MEK1 function. For example, two variants, G128D and G128V that are described in various cancers (39-42) occur in a position, where glycine is 100% conserved. Thus, not only the variants reported so far, but any other substitution of glycine in this position would be damaging. Many disease-associated mutations are found in positions that are not invariable, but still well-conserved. For example, R181M (leads to a loss of a positive charge) variant was described in thyroid carcinoma (43). Our analysis revealed that the only change in position 181 seen during MEK1 evolutionary history is R181K substitution, which maintains the charge and polarity. Thus, any change other than R181K should be classified as damaging.

*De novo* substitution of leucine 42 to phenylalanine was found in an individual with a Cardifaciocutaneous Syndrome (13). Mutation L42F was also found in desmoplastic melanoma sample (44). Position 42 is a part of nuclear export sequence (NES), and according to our MEK1 ortholog MSA (Dataset S2) it endured multiple substitutions from leucine to isoleucine and methionine. Leucine, isoleucine and methionine are all large aliphatic residues. Various organisms were able to tolerate those mutations, what presents strong evidence that corresponding variants are likely to be “benign” in humans. Although phenylalanine is also a large hydrophobic residue, there is no historical evidence to prove that such change could be tolerated, and thus, substitution L42F is predicted to affect the MEK1 biological functioning in a negative way (damaging). Variants Q164K and Q164E were found in patients with CFCS (45) and cancer (46), respectively. Position 164 manifested greater variability. Specifically, glutamine at position 164 had changed multiple times in vertebrates and invertebrates to other polar amino acids such as glutamic and aspartic acids, lysine, arginine, asparagine and serine. Interestingly, all MEK1 proteins from frogs and several snakes contained lysine. Additionally, substitutions to a non-polar small amino acid proline were found in several sequences from different clades and substitutions to hydrophobic leucine, isoleucine and valine were found once each. Thus, hypervariability of this position and the fact that the same substitutions are tolerated in other species suggest that Q164K and Q164E substitutions are likely benign.

Mutation T55P is associated with various RASopathies (45, 47). This position is hypervariable in the MEK1 ortholog set. Multiple changes to other small and large, polar and hydrophobic amino acids in this position occurred throughout MEK1 evolution, However, not a single MEK1 ortholog contains proline in this position. Position T55 is located within N-terminal helix A and T55P substitution may cause a structural damage as prolines are known to cause kinks in alpha helices.

We compared results of our evolutionary analysis of cancer mutations with predictions by popular automated bioinformatics tools PolyPhen-2 (2) and SIFT (3). In four cases, our results disagree and at least in two cases we present additional evidence favoring our interpretation of the variants. Variants R47G, R47Q, P105S and I103S were predicted to be benign by both automated tools (Fig. S3), whereas our analysis suggested that all these variants are damaging, as no such replacements were found throughout the history of MEK1 (Dataset S2).

One of the extensively studied mutations of MEK1, F53L, is predicted as “tolerated” by SIFT. Over the last decade it has been found in multiple cancer tissues (17, 22, 41, 48). F53L has been demonstrated to constitutively activate MEK1, indicated by the increased ERK phosphorylation in cell cultures (17, 49-51), and to severely affect embryonic lethality in *Drosophila* (50, 52). The results of interpretation of F53L mutation using our approach is in agreement with experimental data and is found to be evolutionarily intolerable.

PolyPhen-2 prediction for variant R47G is “benign”, with a score of 0.001. Our analysis suggests that it is “damaging”, because no such substitution can be found in our MEK1 ortholog dataset (Dataset S2). Supporting our interpretation, variant R47G was found in two patients with Langerhans cell histiocytosis (53). Furthermore, position 47 is a part of helix A, formed by residues 43−61. Interaction of helix A with the kinase domain negatively regulates MEK1 kinase activity (54), thus mutations that disrupt this interaction likely result in constitutive ERK pathway activation. Substitutions in positions R47 and adjacent positions (48 and 49) were shown to affect MEK1 function (55). Substitution with alanine led to a 7-fold activation, and insertion of prolines yielded a constitutively active MEK1 (55). The behavior of these substitution mutants was attributed to α-helix disruption (54, 55). Thus, both clinical observation and experimental evidence suggest that R47G mutation is disease-causing. PolyPhen-2 and SIFT predictions for variant R47Q are “benign” (with a score of 0.436) and “tolerated”, respectively. No such substitution can be found in our dataset (Dataset S2) and therefore we interpret it as damaging. Supporting our interpretation, R47Q variant was first detected in breast cancer (invasive ductal carcinoma) cell line (19), and it was later identified in langerhans cell histiocytosis (53, 56). The transforming ability of R47Q was confirmed by using a focus formation assay with mouse 3T3 fibroblasts, and activation potential was evaluated by the increase in the phosphorylation level of direct downstream kinases ERK1/2 (19).

Similarly, our interpretation of variant P105S as “damaging” differs from that by automated tools: “benign” by PolyPhen-2, with a score of 0.418, and “tolerated” by SIFT. This variant was reported in patients with langerhans cell histiocytosis (53), favoring our interpretation.

### Experimental validation of selected variants of unknown significance

To validate our approach experimentally we selected three variants of unknown significance - F53Y, Q56P and G128D - for which there was no clear consensus between ten most widely used automated prediction tools (Fig. 5). The F53Y variant was reported as a rare melanoma mutation (57). The G128D variant was found in Langerhans cell histiocytosis (23). Q56P was originally found in rat fibroblasts as well as in human gastric and lung cancers (19, 58, 59). All three variants were identified as “evolutionary intolerable” and therefore “damaging” by our approach, although their conservation patterns were quite different. While position 128 is 100% invariable, position 53 had one change event to a similar aromatic amino acid, and position 56 is variable.

**Figure 5.**
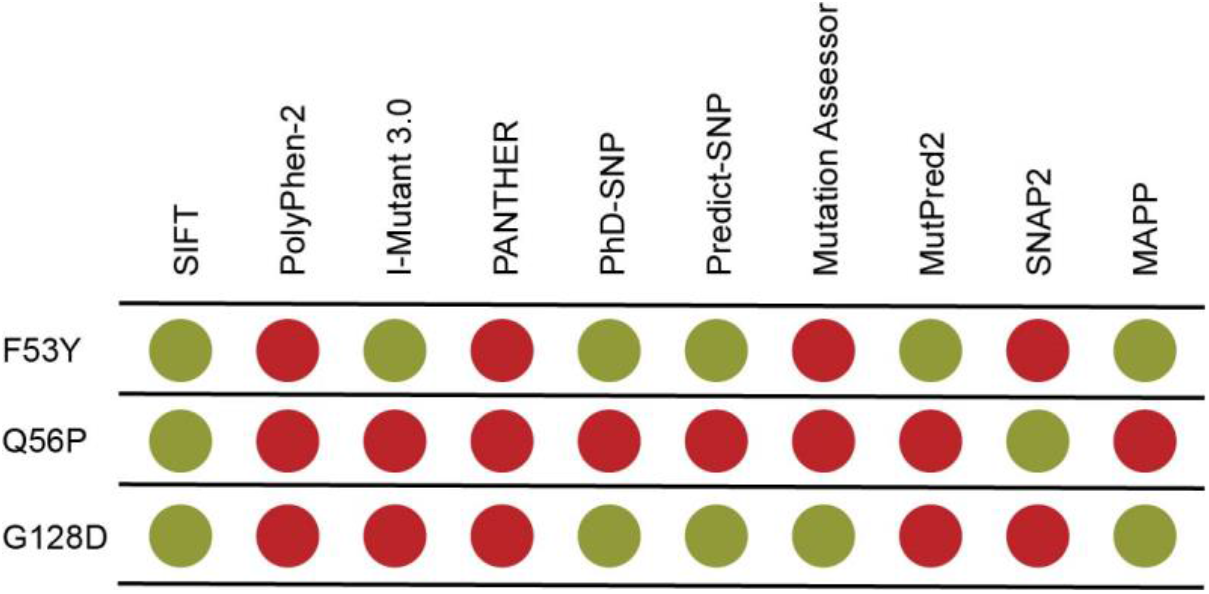
Prediction of functional consequences for selected MEK1 variants of unknown significance by different automated tools (2, 3, 77-84). Green circle represents “benign” and red circle – “pathogenic”.

First, the selected mutations were tested using a misexpression assay in *Drosophila* embryos, as previously described (52). Previously, we found that mutations that cause lethality of *Drosophila* embryos belong to the gain-of-function type mutations (52). Gain-of-function mutations allow for signaling, even in the absence of upstream pathway activation, as well as a more rapid response to activation by RAF even in the absence of necessary adapter proteins (60-62). All three mutations significantly increased embryonic lethality (Fig. 6A), which indicated that all three variants, F53Y, Q56P and G128D, are leading to MEK1 activation.

**Figure 6.**
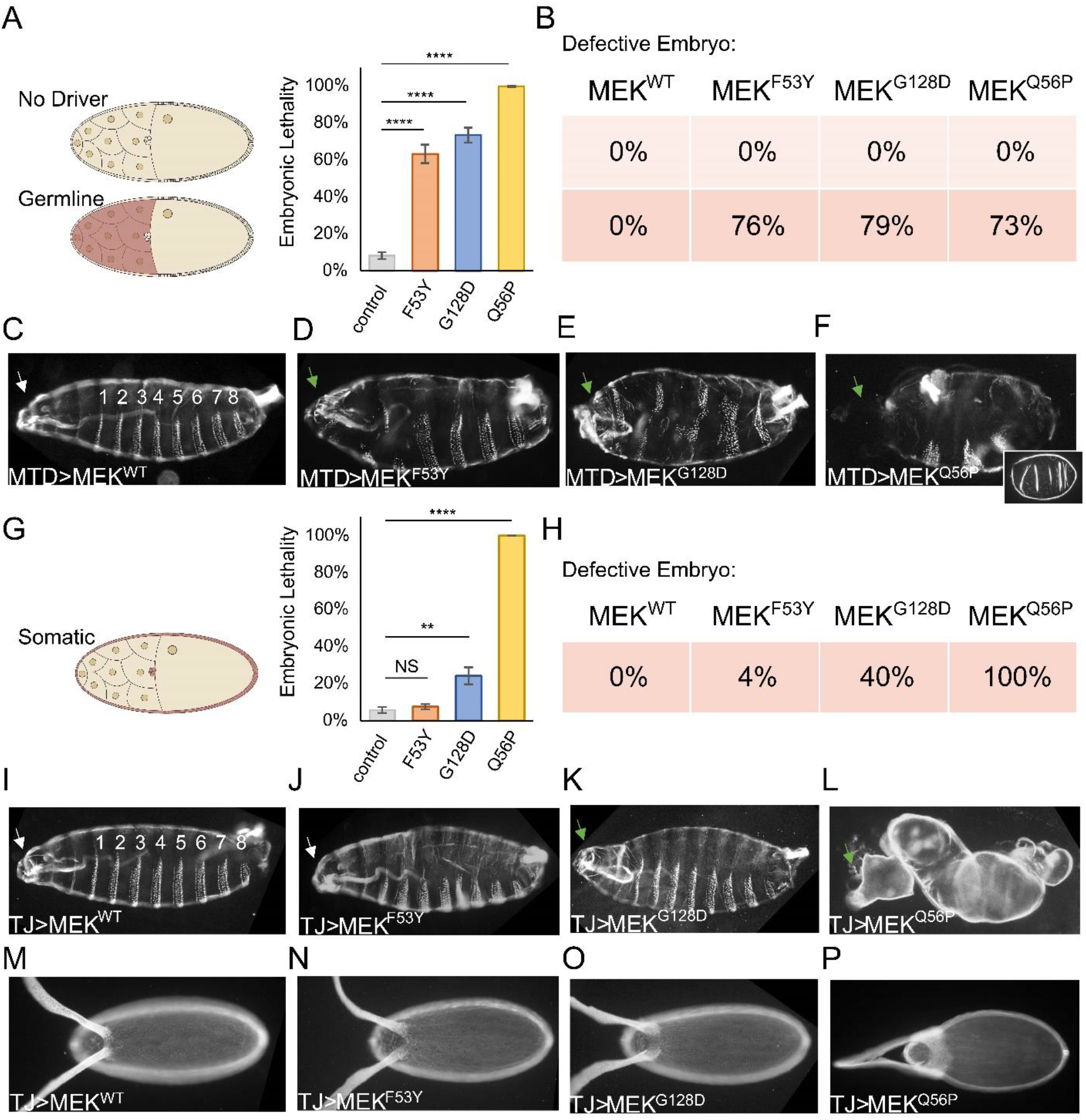
Activating mutations F53Y, G128D, and Q56P in MEK1 cause developmental defects. (A) Lethality of all three mutations is increased when expressed under control of maternal driver (MTD-gal4). The control is overexpression of *mek*^*WT*^ (n=1476). *mek*^*F53Y*^ (n=2596); *mek*^*G128D*^ (n=1742); *mek*^*Q56P*^ (n=727). (B) Defects in embryo patterning are absent without the Gal4 (top). Defects in embryo patterning are frequent in all three mutants, when under control of a maternal germline driver (MTD-Gal4) (bottom, red). (C) Normal embryonic cuticle with the 8 denticle belts shown as well as a normal head (white arrow). *mek*^*WT*^ is phenotypically normal (n=46). (D-F) Representative embryo cuticles show defects in the maternal patterning system. Head pattering is affected by all three mutants (green arrow). (D) *mek*^*F53Y*^ cuticles have disrupted heads as well as fused or missing denticle belts (n=83). (E) *mek*^*G128D*^ cuticles have missing heads as well as fused or missing denticle belts (n=24). (F) *mek*^*Q56P*^ cuticles have missing heads as well as fused or missing denticle belts as well as a failure to retract the germband (n=61). Additionally, 61% of embryos fail to form a cuticle (inset). (G) Lethality of all three mutations is increased when expressed under control of the somatic follicle cell driver (TJ-gal4). The control is overexpression of *mek*^*WT*^ (n=2606). *mek*^*F53Y*^ (n=1167); *mek*^*G128D*^ (n=1351); *mek*^*Q56P*^ (n=223). (H) A graded effect is seen in the embryo cuticles of the three mutants, when under control of a maternal somatic driver (TJ-Gal4) (red). (I) Normally, embryos have 8 segments, visualized by the presence of 8 denticle belts on the ventral side of the embryo. *mek*^*WT*^ is phenotypically normal (n=13). (J-L) Severity increases between the three mutants, which are dorsalized in a graded manor. Dorsalized eggshells result in dorsalized embryos with malformed heads (green arrows), as evidenced in the mutants *mek*^*G128D*^ and *mek*^*Q56P*^. (J) *mek*^*F53Y*^ is phenotypically normal (n=48). (K) *mek*^*G128D*^ is weakly dorsalized (n=43). (L) *mek*^*Q56P*^ is very strongly dorsalized and has no detectable denticle belts (n=30). (M) Normal eggshells show normal spacing of the respiratory filaments (n=18). (N-P) Dorsalized embryos are caused by dorsalized eggshells.
(N) F53Y eggshells are phenotypically normal (n=14). (O) G128D eggshells are weakly dorsailzed, as evidenced by the increase in the spacing between the two respiratory appendages (n=12). (P) Q56P eggshells are more strongly dorsalized (n=16).

Next, we thoroughly examined cuticles of the wild-type embryos and embryos carrying mutated MEK1. Normal embryonic cuticles have a segmented trunk with 8 denticle belts, together with head and tail segments. In the first RAS signaling event of embryogenesis, the activated ligand Trunk is only present in the termini of the embryo, carving out the head and tail segments. We used misexpression by GAL4/UAS to drive mutant MEK1 expression in the early embryo. Mutant embryos without GAL4 had wild type cuticles (Fig. 6B top, C). Embryos with all three mutations in MEK1 developed with erased trunk fates (i.e. denticle belts) in the middle of the cuticle, an area which normally lacks RAS signaling. Additionally, the early occurrence of ectopic signaling was leading to the induction of a negative feedback loop, which reduced signaling in the termini, leading to a loss of head structures (Fig. 6B bottom, D-F). The same phenotype was previously reported for other gain-of function MEK1 mutants (60, 61). Notably, ∼60% of embryos expressing Q56P MEK1 failed to form any cuticle.

Next, we turned to the maternal patterning of the eggshell/egg. This patterning event sets up the dorso-ventral (DV) axis of the eggshell and embryo via EGFR signaling in somatic cells (63-65). The DV patterning of the eggshell is a sensitive system to assay defects in RAS signaling (66). Somatic cells with MEK1 gain-of-function mutants are expected to develop a dorsalized eggshell/embryo. Expression of mutant MEK1 under control of a somatic driver significantly increased embryonic lethality (Fig. 6G). The fraction of abnormal cuticles and lethality agreed with the severity of embryo dorsalization. (Fig. 6H-L). Somatic expression of MEK1 Q56P variant resulted in cuticles that are strongly dorsalized, phenocopying embryos that are null for Dorsal protein (67). The eggshell patterning by these mutants should also be disrupted. Normally, the respiratory tubes on the dorsal side of the eggshell are positioned by an intermediate level of RAS signaling, established by EGFR ligand localized to dorso-anterior and result in a precisely patterned eggshell (Fig. 6M) (66). Increased RAS signaling forces these tubes to be positioned further apart in a graded manor (64). The MEK1 mutants created eggshells that are increasingly dorsalized by mutants of greater strength (Fig. 6N-P). Taken together, the expression of F53Y, Q56P and G128D MEK1 mutants in two different tissues shows that these variants are pathogenic with different level of severity.

## Discussion

The ever-increasing number of genome sequencing studies revealed dozens of variations in MEK1 sequence and many more are likely to be discovered. Only a small portion of those mutations has been proven to affect MEK1 activity, thus leading to developmental diseases and cancers. However, for most of them it is still unclear how and if those variants affect protein function. The functional consequences of the mutations may be established, for example, by testing the kinase phosphorylation activity *in vitro* (49) or analyzing morphological effects in zebrafish or fruit flies (52). These methods present indisputable advantages, and it would be ideal to test all discovered mutations experimentally. Unfortunately, such approaches are not feasible for the analysis of large amounts of genetic variants: *in vitro* and *in vivo* methods are costly and time consuming. Facing these challenges, upon finding new variant clinicians usually turn to automated prediction tools, such as PolyPhen-2 (2) and SIFT (3). Automated sequences searches employed in these tools and other similar methods cannot identify possible duplication events in the genes’ histories, hence multiple sequence alignments that are used for variant interpretation usually include both orthologs and paralogs of the gene of interest, which leads to false negatives.

The presence of seven human MEK proteins increases the chances of including paralogs into the analysis. To avoid this, we have established a precise evolutionary history of the MEK family and identified duplication events. A phylogenetic study of the MEK family conducted in 1999 (32) determined basic relationships between the family members and suggested that its diversification in fungal and animal lineages involved a series of gene duplication events. However, due to a small number of sequences available at that time, this analysis was limited in scope and detailed evolutionary history of this pathway remained unexplored. In the past decade, the number of sequenced eukaryotic genomes increased dramatically, and it is currently not feasible to include all of them in the analysis. Furthermore, despite the availability of a large number of newly sequenced genomes, the distribution of these genomes across different eukaryotic lineages is not uniform: some lineages are well represented, but for many others representation is sparce. Such imbalance could easily lead to a biased perspective on evolutionary history. Therefore, for our analysis we have selected 39 well-assembled high-quality genomes from as many eukaryotic lineages as possible. The resulting phyletic distribution of MEK homologs across these 39 organisms (Fig. 3) agreed with the previous finding that MEK duplications occurred independently in metazoa and fungi (32) and it shows that various independent MEK duplications happened in all other major eukaryotic supergroups. This approach allowed us to establish that only metazoan organisms should be included into the final dataset for variant interpretation.

The obtained MEK1 ortholog MSA was scrutinized from several different angles. The presence of both highly variable and conserved positions showed that the dataset had a sufficient evolutionary depth for an adequate analysis of observed variants. Next, we collected the data on known functionally and structurally important amino acid residues, and found that, satisfactorily, they are conserved in our dataset. Further validation of the dataset against all known experimentally tested disease causing (positive control) and high-frequency MEK1 mutations (negative control), showed that by our analysis they were interpreted as evolutionarily intolerable or tolerable, respectively. This allowed us to conclude that the resulting dataset of MEK1 orthologs was of high-quality and suitable for reliable variant interpretation.

Using our dataset of MEK1 sequences we analyzed all missense mutations in MEK1 reported in COSMIC and ClinVar databases. The majority of found mutations could be confidently assigned as “tolerable” or “intolerable” based on the presence or absence of the variant throughout the master MSA dataset.

Fortunately, we now have enough experimental data on some of the variants that allowed us to validate our dataset. For example, our analysis suggests that mutations C121S, E203K and P124L are intolerable, and there is a compelling evidence that they are indeed alter protein function and embryonic phenotype and therefore are thought to be disease causing (49, 52). However, for the most found protein variations no experimental data available and our analysis provides clues to the outcome of these mutations. For example, I111, L118 and R227 are 100% conserved in our dataset, indicating that any other amino acid in these positions is evolutionary intolerable and thus very likely would be disease causative. Conversely in some cases, such as R96K and S72G, we found evidence that the same changes happened on several occasions, which points out that they are unlikely to be disease-causing.

Upon comparison of our conclusions with predictions made by SIFT and PolyPhen-2 we found a number of conflicting interpretations. For example, several mutations that were assigned as intolerable or uncertain by our approach were predicted to be benign by the automated tools. Surprisingly, even some of the most extensively studied variants, F53L and K57E (49, 52), which are firmly established to cause major MEK1 function alterations, are predicted to be tolerable by SIFT. We analyzed the MEK1 MSA used by PolyPhen-2 and detected not only paralogous sequences (namely MEK2, MEK6 and MEK4), but also unrelated kinases from fungi, that carried inquired variants and thus led to erroneous interpretation.

Based on our analysis we selected F53Y, Q56P and G128D MEK1 mutations to test them experimentally using drosophila embryos. These three mutations were found in different cancer tissues (23, 57, 58) and reported either in COSMIC or ClinVar databases. By our approach each of these mutations was concluded to be “evolutionarily intolerable” and therefore “damaging”, however, automated tools failed to generate a consensus for the outcome (Fig. 5). The missexpression array suggests that all three substitutions lead to the constitutive activation of MEK1 kinase, and, because of the MEK1 activity the number of the lethal or defective embryos increased significantly in each case. Although the severity of each mutation is different, with Q56P being the most severe one, the data obtained from two different patterning systems expressing F53Y, Q56P and G128D MEK1 mutants demonstrated that these variants were all pathogenic.

We could not reliably interpret the evolutionary tolerability for a small set of MEK1 variants (∼13%), due to their rare occurrence in available metazoan genomes. This apparent weakness will be diminished as more genomes become available. On the other hand, the uncertainty of current interpretations for such mutations makes them, rather than reliably predicted variants, a perfect target for experimental validation. Nature had millions of years to experiment with MEK1 protein sequence, and results of many successful experiments are recorded in the genomes of the living species. Here we demonstrate that by careful analysis of these recordings we can improve our assessment of novel variants in MEK1. Our approach should also contribute to improving automated tools for disease-associated variant interpretation.

## Materials and Methods

### Defining evolution of MEK family

To identify closely related proteins, human MEK1 protein sequence (NP_002746.1) was used as query search against the human genome using BLASTP tool. Human MEK1 protein sequence was then queried against a subset of individual genomes of Eukaryote species in NCBI Refseq or non-redundant protein databases (species are listed on Fig. 3). The 10^−35^ E-value cut-off and coverage of more than 75% parameters were used for each of the separate BLASTP searches. All collected sequences were used to generate a multiple sequence alignment (MSA) using MAFFT v7 L-INS-i algorithm (68). The MSA was manually trimmed to include only the kinase domain. Partial and redundant sequences were removed from the alignment. Sequence alignment editing was performed in Jalview (69). We used two approaches for phylogenetic relationship reconstruction, maximum likelihood and neighbor joining methods with 1000 bootstrap replications. Phylogenetic inference was conducted using with MEGAX package (70). Phylogenetic trees were analyzed and visualized using iTOL tool (71).

### Orthology assignment and MEK1 orthologs master set

Orthologs and paralogs were distinguished using several approaches: (i) by identifying reciprocal best BLAST hits between protein sequences of two organisms; (ii) domain architecture of the group of proteins; (iii) and confirmed by the presence of monophyletic clades of each group of the orthologs in the phylogenetic trees.

For the broader panel of MEK1 sequences human MEK1 protein sequence was used to query the RefSeq database of metazoan species using BLAST. Retrieved sequences were aligned with MAFFT v7.154b E-INS-i (68). MEK2, MEK5, MEK3/6 and MEK4/7 sequences from representative species set, assigned earlier, were added to the analysis as outgroups. Neighbor joining and maximum likelihood phylogenetic trees were generated with 500 bootstrap replications with MEGAX (70). The outgroups on the tree were not considered to be MEK1 true orthologs and were discarded from the master MSA of MEK1 ortholog sequences set.

### Interpretation of variants

ClinVar, COSMIC (72) and literature were searched to identify missense mutations that were reported to be associated with cancer or any type of RASopathy. The Genome Aggregation Database (gnomAD, (38)) database was searched to identify MEK1 variants that occur frequently in human populations.

To assign the significance of each of the mutations we used SAVER algorithm (6). Briefly, this approach focuses on (i) making a set of sequences containing only functional orthologs, which requires the establishment of evolutionary history of a protein, and (ii) counting how many times the substitution occurred independently during evolutionary history of the protein, rather than utilizing the abundance of the variant in MSA, to neutralize bias in the alignment created by the inclusion of sequences that are very closely related to each other.

### Experimental procedures

*Fly stocks*. Fly stocks were maintained under standard conditions and crosses were performed at 25°C. MTD-GAL4, UAS-MEKWT, and TJ-GAL4 flies were described previously (52, 73, 74). cDNA for Dsor1, the Drosophila homologue of MEK, was cloned into an intermediate vector, pMiniT, and mutated by PCR to generate the disease variant alleles (New England Biolabs #E0554S). The utilized mutagenic primers are as follows: F53S-ATCAAGATGTaCCTCAGCCAGAAGG, G128D-CACATTGTCGaTTTCTACGGC, and Q56P-TTCCTCAGCCcGAAGGAGAAG. The UAS-MEK variants were cloned into pTIGER and integrated into attP40 site using the ΦC31-based integration system as described previously (60).

*Cuticle phenotyping*. Embryos were dechorionated with 50% bleach after being aged for 22 h as previously reported (61, 75). Dechorionated embryos were shaken in methanol and heptane (1:1) and incubated overnight in a media containing lactic acid and Hoyer’s media (1:1) at 65°C. Embryos were imaged in darkfield on a Nikon Eclipse Ni.

*Embryonic lethality*. Lethality was determined by setting up egg lay cages for a period allowing for ∼300 eggs to be oviposited onto grape juice agar. Plates were aged for greater than 30 h and lethality was determined as the fraction of unhatched eggs. Replicates were conducted to allow for survey of over 1000 eggs for each genotype.

## Acknowledgments

This work was supported by NIH grants R35-GM131760 (to I.B.Z.) and R01-GM086537 and R01-HD085870 (to S.Y.S.).

