## Supplementary Figures and Tables for "Evolutionary history of MEK1 illuminates the nature of cancer and RASopathy mutations"

**This PDF file includes:**

Figures S1 to S2
Tables S1 to S2
Legends for Datasets S1 to S3
SI References

**Other supplementary materials for this manuscript include the following:**

Datasets S1 to S3

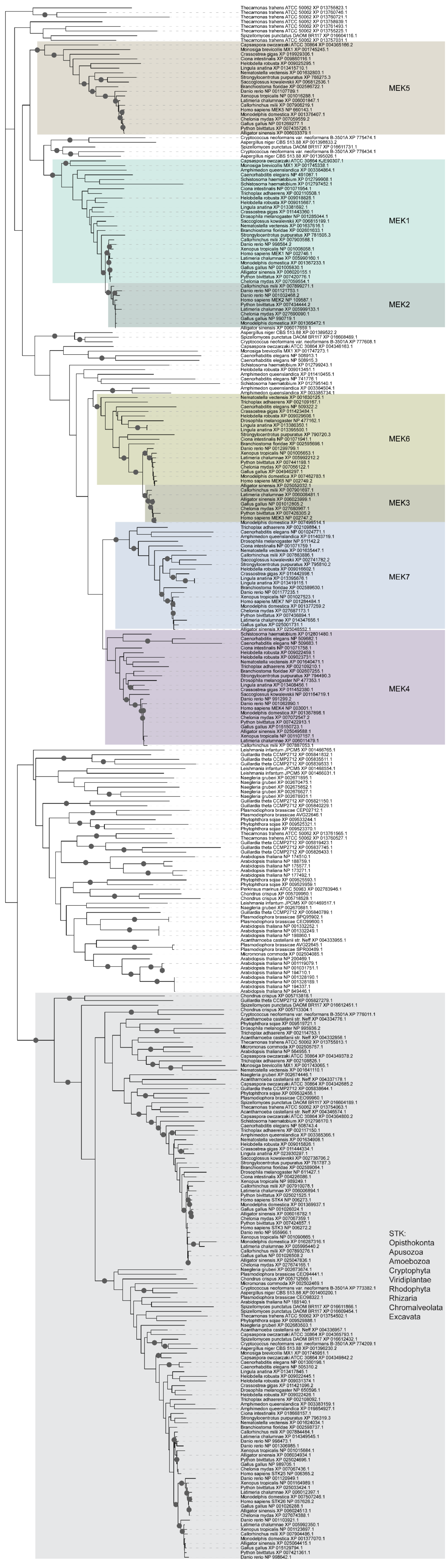


**Fig. S1. Maximum likelihood phylogenetic tree of the MEK protein family and their closest homologs from the representative set of eukaryotic species.** Bootstrap support values >50% are indicated as circles.

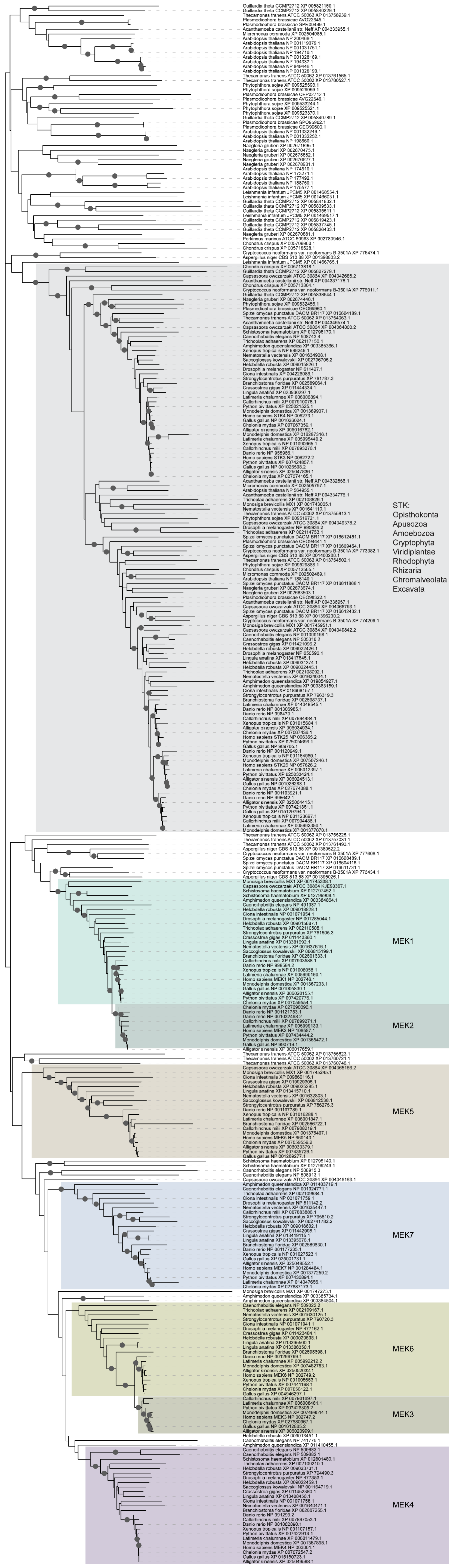


**Fig. S2. Neighbor joining phylogenetic tree of the MEK protein family and their closest homologs from the representative set of eukaryotic species.** Bootstrap support values >50% are indicated as circles

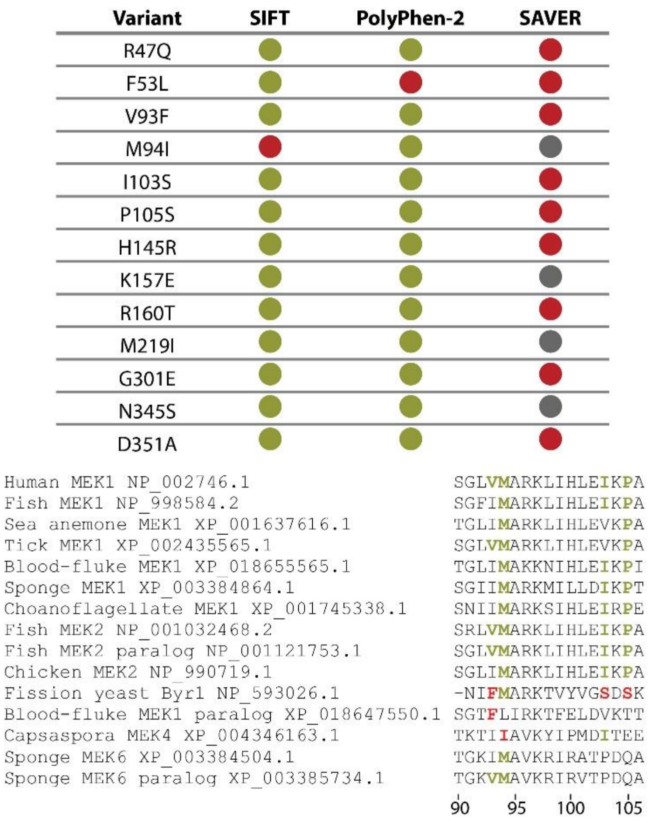


**Fig. S3. Inclusion of paralogs affects predictions of missense mutations outcome by automated tools.** Top: Comparison of predicting the missense mutation effect by different methods. Red circle represents “predicted as damaging”, the green circle represents “predicted as benign”, grey circle represents “uncertain” in our analysis. Bottom: Examples of protein sequences erroneously included into MEK1 analysis by PolyPhen-2. Multiple inclusions of paralogous sequences were identified, such as MEK2, MEK6, MEK4 and distinct kinases from fungi. Residues highlighted in red in these sequences are the potential causes of erroneous predictions. Numeration is according to the human MEK1 protein sequence.

.

**Table S1. Conservation of the functionally assigned positions of human MEK1.** Based on (1-11)

| Position in human MEK1 | Amino acid in human MEK | Function | Conservation in MEK family and homologs from the representative set of eukaryote (Dataset S1) | Conservation in MEK1 orthologs from Metazoa (Dataset S2) |
|---|---|---|---|---|
| 75 | G | ATP-phosphate binding loop, P-loop | invariable | invariable |
| 77 | G | ATP-phosphate binding loop, P-loop | 1 sequence has R | invariable |
| 78 | N | Interaction with MEK2 | | 1 sequence has S |
| 80 | G | ATP-phosphate binding loop, P-loop | invariable | invariable |
| 81 | V | Catalytic spine | | invariable |
| 95 | A | Catalytic spine | | invariable |
| 97 | K | Catalytic core | invariable | invariable |
| 114 | E | Activation core | negatively charged | invariable |
| 118 | L | Regulatory spine | | invariable |
| 129 | F | Regulatory spine | | |
| 138 | E | Interaction with BRAF | | |
| 144 | E | ATP binding | | 1 sequence has Q |
| 146 | M | ATP binding | | invariable |
| 151 | L | Catalytic spine | | 1 sequence has F |
| 188 | H | Regulatory spine | invariable | invariable |
| 189 | R | Catalytic loop | invariable | invariable |
| 190 | D | Catalytic core | | invariable |
| 192 | K | Catalytic loop | invariable | 1 sequence has Q |
| 196 | I | Catalytic spine | | |
| 197 | L | Catalytic spine | | invariable |
| 198 | V | Catalytic spine | | |
| 208 | D | Catalytic core | invariable | invariable |
| 209 | F | Regulatory spine | | invariable |
| 210 | G | Activation segment | | invariable |
| 212 | S | phosphorylation site | | invariable |
| 218 | S | RAF phosphorylation site | | invariable |
| 220 | A | Interaction with KSR2 | | invariable |
| 221 | N | Interaction with KSR2 | | |
| 222 | S | RAF phosphorylation site | | |
| 224 | V | Interaction with KSR2 | | invariable |
| 225 | G | Interaction with KSR2 | 1 sequence has E | invariable |
| 226 | T | Interaction with KSR2 | | 1 event of a change to S |
| 228 | S | Interaction with BRAF | | invariable |
| 230 | M | Interaction with KSR2 and BRAF | | invariable |
| 235 | L | Interaction with KSR2 | | |
| 245 | D | Structural role | negatively charged | invariable |

| | | | | |
|---|---|---|---|---|
| 253 | L | Catalytic spine | | |
| 256 | M | Catalytic spine | | |
| 286 | T | Cdk5 phosphorylation site | | |
| 292 | T | ERK phosphorylation site | | |
| 298 | S | PAK1 phosphorylation site | | |
| 308 | M | Interaction with KSR1 | | |
| 309 | A | Interaction with KSR2 | | |
| 310 | I | Interaction with BRAF and KSR1 | | |
| 311 | F | Interaction with KSR2 | | invariable |
| 314 | L | Interaction with BRAF | | invariable |
| 315 | D | Interaction with KSR2 | | |
| 318 | V | Interaction with KSR2 | | |
| 386 | T | Cdk5 and ERK phosphorylation site | | |

**Table S2. MEK1 variations found in COSMIC (12) and ClinVar databases and their interpretation by PolyPhen-2 (13), SIFT (14) and SAVER (15) algorithms.** For mutation found in proline-rich segment (indicated by asterisk) evolutionarily tolerability assessment is based on MEK1 orthologs from Gnathostomata species.

| Variant | Observation | Evolutionary permissibility | PolyPhen-2 prediction | SIFT prediction |
|---|---|---|---|---|
| L42F | Phe is not present in any Metazoan species | Not permitted | benign | affect protein function |
| L42H | His is not present in any Metazoan species | Not permitted | probably damaging | affect protein function |
| E44K | Lys is not present in any Metazoan species | Not permitted | possibly damaging | affect protein function |
| E44G | Gly is not present in any Metazoan species | Not permitted | probably damaging | affect protein function |
| Q46L | Leu is not present in any Metazoan species | Not permitted | benign | affect protein function |
| R47Q | Gln is not present in any Metazoan species | Not permitted | possibly damaging | tolerated |
| R49C | Cys is not present in any Metazoan species | Not permitted | probably damaging | affect protein function |
| R49H | His is not present in any Metazoan species | Not permitted | probably damaging | affect protein function |
| R49L | Leu is not present in any Metazoan species | Not permitted | probably damaging | affect protein function |
| L50V | Val is not present in any Metazoan species | Not permitted | possibly damaging | tolerated |
| L50P | Pro is not present in any Metazoan species | Not permitted | possibly damaging | affect protein function |
| E51G | Gly is not present in any Metazoan species | Not permitted | probably damaging | tolerated |
| F53I | Ile is not present in any Metazoan species | Not permitted | probably damaging | affect protein function |
| F53V | Val is not present in any Metazoan species | Not permitted | probably damaging | affect protein function |
| F53L | Leu is not present in any Metazoan species | Not permitted | probably damaging | tolerated |
| F53S | Ser is not present in any Metazoan species | Not permitted | probably damaging | affect protein function |
| F53Y | Tyr is not present in any Metazoan species | Not permitted | probably damaging | tolerated |
| F53C | Cys is not present in any Metazoan species | Not permitted | probably damaging | affect protein function |
| L54P | Pro is not present in any Metazoan species | Not permitted | probably damaging | affect protein function |
| T55P | Pro is not present in any Metazoan species | Not permitted | probably damaging | tolerated |
| Q56P | Pro is not present in any Metazoan species | Not permitted | probably damaging | tolerated |
| Q56K | 2 events of change to Lys | Permitted | benign | tolerated |
| Q56H | His is not present in any Metazoan species | Not permitted | possibly damaging | tolerated |
| K57E | 100% Lys | Not permitted | probably damaging | tolerated |

| | | | | |
|---|---|---|---|---|
| K57Q | 100% Lys | Not permitted | probably damaging | tolerated |
| K57T | 100% Lys | Not permitted | probably damaging | tolerated |
| K57N | 100% Lys | Not permitted | probably damaging | tolerated |
| K57R | 100% Lys | Not permitted | probably damaging | tolerated |
| Q58H | Only one sequence with His | Uncertain | benign | tolerated |
| V60G | Gly is not present in any Metazoan species | Not permitted | probably damaging | affect protein function |
| V60E | Glu is not present in any Metazoan species | Not permitted | probably damaging | affect protein function |
| V60M | Only one sequence with Met. Ile and Leu >2 events | Uncertain | benign | affect protein function |
| K64E | Glu is not present in any Metazoan species | Not permitted | benign | tolerated |
| D67N | Only one sequence with Asn | Uncertain | possibly damaging | tolerated |
| D67Y | Tyr is not present in any Metazoan species | Not permitted | probably damaging | affect protein function |
| E69G | Gly is not present in any Metazoan species | Not permitted | probably damaging | affect protein function |
| E69K | Only one sequence with Lys | Uncertain | benign | tolerated |
| S72G | >2 events of change to Gly | Permitted | benign | tolerated |
| A76S | >2 events of change to Ser | Permitted | benign | tolerated |
| A76V | Val is not present in any Metazoan species | Not permitted | probably damaging | tolerated |
| G77D | 100% Gly | Not permitted | probably damaging | affect protein function |
| N78S | Only one sequence with Ser. No changes otherwise. | Not permitted | possibly damaging | affect protein function |
| G79S | Only one sequence with Ser. 1 event of change to Trp. Nothing else. | Not permitted | probably damaging | tolerated |
| G79V | Val is not present in any Metazoan species | Not permitted | probably damaging | affect protein function |
| G80S | 100% Gly | Not permitted | probably damaging | affect protein function |
| V82M | 100% Val | Not permitted | probably damaging | affect protein function |
| K84R | >2 events of change to Arg | Permitted | benign | tolerated |
| S86A | 2 events of change to Ala | Permitted | benign | tolerated |
| K88R | >2 events of change to Arg | Permitted | benign | tolerated |
| P89T | One event of change to Thr | Uncertain | benign | affect protein function |
| L92R | Arg is not present in any Metazoan species | Not permitted | possibly damaging | affect protein function |
| V93I | >2 events of change to Ile | Permitted | benign | tolerated |
| V93F | Phe is not present in any Metazoan species | Not permitted | benign | tolerated |
| M94I | Only one sequence with Ile. No other changes | Not permitted | benign | affect protein function |
| R96K | >2 events of change to Lys | Permitted | possibly damaging | tolerated |

| | | | | |
|---|---|---|---|---|
| I99T | Thr is not present in any Metazoan species | Not permitted | probably damaging | affect protein function |
| H100N | Asn is not present in any Metazoan species | Not permitted | probably damaging | tolerated |
| E102G | Gly is not present in any Metazoan species. Only negatively charged | Not permitted | possibly damaging | tolerated |
| I103N | Asn is not present in any Metazoan species. Val and Ile only | Not permitted | probably damaging | affect protein function |
| I103S | Ser is not present in any Metazoan species. Val and Ile only | Not permitted | possibly damaging | tolerated |
| I103M | Met is not present in any Metazoan species. Val and Ile only | Not permitted | probably damaging | affect protein function |
| A106T | Thr 2 events | Permitted | benign | tolerated |
| R108W | Only one sequence with Trp. | Uncertain | probably damaging | affect protein function |
| R108L | Only one sequence with Leu | Uncertain | probably damaging | tolerated |
| R108Q | Gln is not present in any Metazoan species | Not permitted | probably damaging | tolerated |
| I111N | 100% Ile | Not permitted | probably damaging | affect protein function |
| I111S | 100% Ile | Not permitted | probably damaging | affect protein function |
| L115V | 100% Leu | Not permitted | probably damaging | affect protein function |
| L115P | 100% Leu | Not permitted | probably damaging | affect protein function |
| L115V | 100% Leu | Not permitted | probably damaging | affect protein function |
| L118V | 100% Leu | Not permitted | probably damaging | affect protein function |
| H119Y | Tyr is not present in any Metazoan species | Not permitted | probably damaging | tolerated |
| H119Q | Gln is not present in any Metazoan species | Not permitted | probably damaging | affect protein function |
| H119P | Pro is not present in any Metazoan species | Not permitted | probably damaging | affect protein function |
| E120D | >2 events of change to Asp | Permitted | benign | tolerated |
| C121R | 100% Cys | Not permitted | probably damaging | affect protein function |
| C121S | 100% Cys | Not permitted | probably damaging | affect protein function |
| C121G | 100% Cys | Not permitted | probably damaging | affect protein function |
| N122D | Asp is not present in any Metazoan species | Not permitted | probably damaging | tolerated |
| S123P | Pro is not present in any Metazoan species | Not permitted | probably damaging | affect protein function |
| P124S | Only one sequence with Ser. Conservation for small residues. | Uncertain | probably damaging | tolerated |
| P124T | Only one sequence with Thr. Conservation for small residues. | Uncertain | probably damaging | affect protein function |
| P124R | Arg is not present in any Metazoan species | Not permitted | probably damaging | affect protein function |
| P124L | Leu is not present in any Metazoan species | Not permitted | probably damaging | affect protein function |
| P124Q | Gln is not present in any Metazoan species | Not permitted | probably damaging | affect protein function |

| | | | | |
|---|---|---|---|---|
| P124M | Met is not present in any Metazoan species. Conservation for small residues. | Not permitted | probably damaging | affect protein function |
| Y125C | Cys is not present in any Metazoan species. Aromatic residues | Not permitted | probably damaging | affect protein function |
| V127M | Met is not present in any Metazoan species. Val and 1 event of change to Ile | Not permitted | probably damaging | affect protein function |
| G128A | 100% Gly | Not permitted | probably damaging | affect protein function |
| G128D | 100% Gly | Not permitted | probably damaging | tolerated |
| G128V | 100% Gly | Not permitted | probably damaging | affect protein function |
| G128C | 100% Gly | Not permitted | probably damaging | affect protein function |
| F129L | Leu is not present in any Metazoan species | Not permitted | probably damaging | affect protein function |
| Y130N | Asn is not present in any Metazoan species | Not permitted | probably damaging | affect protein function |
| Y130H | His is not present in any Metazoan species | Not permitted | probably damaging | affect protein function |
| Y130C | Cys is not present in any Metazoan species | Not permitted | probably damaging | affect protein function |
| A132V | Only one sequence with Val | Uncertain | probably damaging | affect protein function |
| Y134C | One event of change to Cys | Uncertain | probably damaging | affect protein function |
| D136N | Asn 2 events | Permitted | benign | tolerated |
| D136Y | Tyr is not present in any Metazoan species | Not permitted | probably damaging | affect protein function |
| G137V | Val is not present in any Metazoan species | Not permitted | probably damaging | affect protein function |
| E138K | Lys is not present in any Metazoan species | Not permitted | possibly damaging | tolerated |
| M143V | Val is not present in any Metazoan species | Not permitted | probably damaging | affect protein function |
| M146I | 100% Met | Not permitted | probably damaging | affect protein function |
| S150F | 100% Ser | Not permitted | probably damaging | affect protein function |
| D152N | 100% Asp | Not permitted | probably damaging | affect protein function |
| V154I | >2 events of change to Ile | Permitted | benign | tolerated |
| A158V | One event of change to Val | Uncertain | possibly damaging | tolerated |
| R160T | Thr is not present in any Metazoan species | Not permitted | benign | tolerated |
| I161V | Only one sequence with Val | Uncertain | possibly damaging | tolerated |
| I161T | Thr is not present in any Metazoan species | Not permitted | probably damaging | affect protein function |
| P162S | Ser is not present in any Metazoan species | Not permitted | probably damaging | tolerated |
| P162H | His 2 events | Permitted | possibly damaging | affect protein function |
| Q164E | >2 events of change to Glu | Permitted | benign | tolerated |
| Q164K | >2 events of change to Lys | Permitted | benign | tolerated |

| | | | | |
|---|---|---|---|---|
| K168R | >2 events of change to Arg | Permitted | benign | tolerated |
| A172V | Val is not present in any Metazoan species | Not permitted | probably damaging | affect protein function |
| I174V | >2 events of change to Val | Permitted | benign | tolerated |
| G176S | 100% Gly | Not permitted | possibly damaging | affect protein function |
| G176V | 100% Gly | Not permitted | probably damaging | affect protein function |
| L177M | 100% Leu | Not permitted | probably damaging | affect protein function |
| L177V | 100% Leu | Not permitted | probably damaging | affect protein function |
| R181M | Met is not present in any Metazoan species | Uncertain | probably damaging | tolerated |
| H184R | Only one sequence with Arg | Uncertain | benign | tolerated |
| K185N | >2 events of change to Asn | Permitted | benign | tolerated |
| M187I | >2 events of change to Ile | Permitted | benign | tolerated |
| H188Y | 100% His | Not permitted | probably damaging | affect protein function |
| R189G | 100% Arg | Not permitted | possibly damaging | affect protein function |
| K192N | Asn is not present in any Metazoan species | Not permitted | probably damaging | affect protein function |
| P193S | 100% Pro | Not permitted | probably damaging | affect protein function |
| P193A | 100% Pro | Not permitted | probably damaging | affect protein function |
| S194A | Catalytic loop, Ser 100% | Not permitted | probably damaging | affect protein function |
| I196V | >2 events of change to Val | Permitted | probably damaging | tolerated |
| S200Y | Tyr is not present in any Metazoan species | Not permitted | probably damaging | affect protein function |
| R201S | >2 events of change to Ser | Permitted | probably damaging | tolerated |
| R201C | Cys is not present in any Metazoan species | Not permitted | probably damaging | affect protein function |
| R201H | His 2 events | Permitted | possibly damaging | tolerated |
| E203Q | Gln is not present in any Metazoan species | Not permitted | probably damaging | tolerated |
| E203K | Lys is not present in any Metazoan species | Not permitted | probably damaging | affect protein function |
| E203A | Ala is not present in any Metazoan species | Not permitted | probably damaging | affect protein function |
| E203V | Val is not present in any Metazoan species | Not permitted | probably damaging | affect protein function |
| E203D | One event of change to Asp | Uncertain | possibly damaging | tolerated |
| I204F | Phe is not present in any Metazoan species | Not permitted | probably damaging | affect protein function |
| I204T | Only one sequence with Thr | Uncertain | probably damaging | affect protein function |
| L206F | Phe is not present in any Metazoan species | Not permitted | probably damaging | affect protein function |
| G210R | 100% Gly | Not permitted | probably damaging | affect protein function |

| | | | | |
|---|---|---|---|---|
| V211I | Ile is not present in any Metazoan species | Not permitted | possibly damaging | affect protein function |
| V211D | Asp is not present in any Metazoan species | Not permitted | probably damaging | affect protein function |
| S212N | 100% Ser | Not permitted | probably damaging | affect protein function |
| G213V | Activation segment, 100% Gly | Not permitted | probably damaging | affect protein function |
| L215P | 100% Leu | Not permitted | probably damaging | affect protein function |
| L215I | 100% Leu | Not permitted | probably damaging | affect protein function |
| L215F | 100% Leu | Not permitted | probably damaging | affect protein function |
| S218F | 100% Ser | Not permitted | probably damaging | affect protein function |
| M219I | >2 events of change to Ile | Permitted | benign | tolerated |
| S222Y | Tyr is not present in any Metazoan species | Not permitted | probably damaging | affect protein function |
| V224M | 100% Met | Not permitted | probably damaging | affect protein function |
| R227K | 100% Arg | Not permitted | probably damaging | tolerated |
| Y229H | 100% Tyr | Not permitted | probably damaging | affect protein function |
| S231L | Leu is not present in any Metazoan species | Not permitted | probably damaging | affect protein function |
| L235H | His is not present in any Metazoan species | Not permitted | probably damaging | affect protein function |
| S241Y | Tyr is not present in any Metazoan species | Not permitted | probably damaging | affect protein function |
| V242M | Met is not present in any Metazoan species | Not permitted | probably damaging | affect protein function |
| Q243H | Only one event of change to His | Uncertain | benign | tolerated |
| S244A | Ala is not present in any Metazoan species | Not permitted | possibly damaging | tolerated |
| D245N | 100% Asp | Not permitted | probably damaging | affect protein function |
| I246L | Only one sequence with Leu | Uncertain | benign | affect protein function |
| W247R | Arg is not present in any Metazoan species | Not permitted | probably damaging | affect protein function |
| E255K | 100% Glu | Not permitted | probably damaging | affect protein function |
| M256L | >2 events of change to Leu | Permitted | benign | tolerated |
| M256T | Thr is not present in any Metazoan species | Not permitted | possibly damaging | affect protein function |
| A257V | Only one sequence with Val | Uncertain | possibly damaging | affect protein function |
| V258I | >2 events of change to Ile | Permitted | benign | tolerated |
| P264S* | 100% Pro in Gnathostomata species | Not permitted | probably damaging | affect protein function |
| P264H* | 100% Pro in Gnathostomata species | Not permitted | probably damaging | affect protein function |
| P264R* | 100% Pro in Gnathostomata species | Not permitted | probably damaging | affect protein function |
| P265S* | Only one sequence with Ser among Gnathostomata species | Uncertain | benign | tolerated |

| | | | | |
|---|---|---|---|---|
| P265L* | Leu is not present in any Metazoan species | Not permitted | possibly damaging | affect protein function |
| A268T* | >2 events of change to Thr | Permitted | benign | tolerated |
| A268V* | Only one sequence with Val among Gnathostomata species | Uncertain | benign | tolerated |
| L271R* | 100% Leu in Gnathostomata species | Not permitted | possibly damaging | tolerated |
| G276W* | Trp is not present in any Gnathostomata species | Not permitted | probably damaging | affect protein function |
| G276R* | Arg is not present in any Metazoan species | Not permitted | benign | affect protein function |
| Q278H* | >2 events of change to His in Gnathostomata species | Permitted | benign | tolerated |
| A283V* | >2 events of change to Val | Permitted | benign | tolerated |
| E285K* | Lys is not present in any Metazoan species | Not permitted | benign | tolerated |
| A283V* | >2 events of change to Val | Permitted | benign | tolerated |
| P288H* | His is not present in any Gnathostomata species | Not permitted | possibly damaging | affect protein function |
| P288L* | Leu is not present in any Metazoan species | Not permitted | benign | affect protein function |
| R291G* | Only one sequence with Gly among Gnathostomata species | Uncertain | benign | tolerated |
| R291K* | Lys is not present in any Gnathostomata species | Not permitted | benign | tolerated |
| T292I* | Ile is not present in any Gnathostomata species | Not permitted | benign | tolerated |
| T292S* | Only one sequence with Ser among Gnathostomata species | Uncertain | benign | tolerated |
| P293S* | Ser is not present in any Gnathostomata species | Not permitted | benign | affect protein function |
| G294R* | 100% Gly in Gnathostomata species | Not permitted | possibly damaging | tolerated |
| G294W* | 100% Gly in Gnathostomata species | Not permitted | probably damaging | affect protein function |
| G294E* | 100% Gly in Gnathostomata species | Not permitted | possibly damaging | tolerated |
| G294V* | 100% Gly in Gnathostomata species | Not permitted | benign | tolerated |
| P296H* | His is not present in any Gnathostomata species | Not permitted | probably damaging | affect protein function |
| G301E* | Glu is not present in any Gnathostomata species | Not permitted | benign | tolerated |
| G301R* | Arg is not present in any Gnathostomata species | Not permitted | probably damaging | tolerated |
| M302L* | >2 events of change to Leu | Permitted | benign | tolerated |
| D303N* | One event of change to Asn | Uncertain | benign | tolerated |
| R305Q* | 100% Arg in Gnathostomata species | Not permitted | possibly damaging | tolerated |
| P306L* | 100% Pro in Gnathostomata species | Not permitted | benign | tolerated |
| P306H* | 100% Pro in Gnathostomata species | Not permitted | possibly damaging | tolerated |
| M308L | >2 events of change to Leu | Permitted | benign | tolerated |
| M308R | Arg is not present in any Metazoan species | Not permitted | possibly damaging | affect protein function |

| | | | | |
|---|---|---|---|---|
| I310L | Leu is not present in any Metazoan species | Not permitted | possibly damaging | affect protein function |
| I310H | His is not present in any Metazoan species | Not permitted | probably damaging | affect protein function |
| E312Q | 100% Glu | Not permitted | probably damaging | tolerated |
| L313F | Phe is not present in any Metazoan species | Not permitted | probably damaging | affect protein function |
| P321S | One event of change to Ser in two genomes of lancelets. Ala >2 events of change to , one sequence with His, Thr, Glu, Gln. | Uncertain | possibly damaging | tolerated |
| P321H | Only one sequence with His | Uncertain | benign | affect protein function |
| P322S | Ser is not present in any Metazoan species | Not permitted | probably damaging | tolerated |
| P323A | Only one sequence with change (Ala) | Not permitted | probably damaging | affect protein function |
| K324Q | >2 events of change to Gln | Permitted | benign | tolerated |
| P326H | His is not present in any Metazoan species | Not permitted | probably damaging | affect protein function |
| S327T | >2 events of change to Thr | Permitted | benign | tolerated |
| G328E | Glu is not present in any Metazoan species | Not permitted | benign | tolerated |
| S331R | Arg is not present in any Metazoan species | Not permitted | possibly damaging | affect protein function |
| L332V | Only one sequence with Val | Uncertain | benign | tolerated |
| L332R | Arg is not present in any Metazoan species | Not permitted | benign | tolerated |
| E333Q | Only one sequence with Gln | Uncertain | benign | tolerated |
| D336H | His is not present in any Metazoan species | Not permitted | probably damaging | tolerated |
| N339I | Ile is not present in any Metazoan species | Not permitted | possibly damaging | affect protein function |
| N345T | Thr is not present in any Metazoan species | Not permitted | possibly damaging | tolerated |
| A347T | >2 events of change to Thr | Permitted | benign | tolerated |
| R349S | 100% Arg | Not permitted | probably damaging | affect protein function |
| R349K | 100% Arg | Not permitted | probably damaging | affect protein function |
| D351A | Ala is not present in any Metazoan species | Not permitted | possibly damaging | tolerated |
| D351G | Only one sequence with Gly | Uncertain | possibly damaging | tolerated |
| D351E | Only one sequence with Glu | Uncertain | benign | tolerated |
| L352F | One event of change to Phe | Uncertain | probably damaging | tolerated |
| Q354H | Only one sequence with His, Phe, Lys, Leu, Gly, Ala. One event of change to Tyr and Glu. More than 2 events of change to Ser, Val, Asn, Thr, Met | Uncertain | benign | tolerated |
| M356V | >2 events of change to Val | Permitted | benign | tolerated |
| H358Y | Only one sequence with Tyr | Uncertain | probably damaging | affect protein function |
| H358Q | Only one sequence with Gln | Uncertain | probably damaging | tolerated |

| | | | | |
|---|---|---|---|---|
| R363I | 2 events of change to Ile | Permitted | possibly damaging | tolerated |
| E367K | >2 events of change to Lys | Permitted | benign | tolerated |
| E367Q | 2 events of change to Gln | Permitted | benign | tolerated |
| E367G | Gly is not present in any Metazoan species | Not permitted | possibly damaging | tolerated |
| V369A | >2 events of change to Ala | Permitted | possibly damaging | tolerated |
| A372V | Only one sequence with Val | Uncertain | probably damaging | affect protein function |
| L375I | >2 events of change to Ile | Permitted | benign | tolerated |
| L375P | Pro is not present in any Metazoan species | Not permitted | probably damaging | affect protein function |
| C376R | Arg 2 events | Permitted | benign | tolerated |
| S377F | Phe is not present in any Metazoan species | Not permitted | possibly damaging | affect protein function |
| I379V | Val 2 events | Permitted | benign | tolerated |
| G380S | Only one sequence with Ser | Uncertain | benign | tolerated |
| L381P | Pro is not present in any Metazoan species | Not permitted | probably damaging | affect protein function |
| N382H | >2 events of change to His | Permitted | benign | tolerated |
| P387S | >2 events of change to Ser | Permitted | possibly damaging | affect protein function |
| A390T | >2 events of change to Thr | Permitted | benign | tolerated |
| A391V | >2 events of change to Val | Permitted | benign | affect protein function |
| V393I | >2 events of change to Ile | Permitted | benign | affect protein function |

**Dataset S1. Protein sequences of MEK family members and their closest relatives from representative species of Eukaryota.**

**Dataset S2. List of MEK1 orthologs from metazoa.**

**Dataset S3. Conservation analysis of MEK1 protein sequence.**
